# Spectral acoustic contributions to musical pleasure are dynamically shaped by autonomic neural inputs

**DOI:** 10.1101/2025.01.05.631396

**Authors:** Vincent K.M. Cheung, Tamaka Harada, Shu Sakamoto, Shinichi Furuya

## Abstract

Most people enjoy music and often use music to regulate their emotions. Although pleasure derived from music-listening has been shown to be mediated by dopaminergic signals in the mesolimbic reward network, its relationship with physiology is still poorly understood. Here, we introduced *time-warped representational similarity analysis* (twRSA) to directly map dynamic representations of multiple modalities across variable-duration stimuli. Our method revealed that although time-varying spectral and tonal acoustic features predicted changes in autonomic neural responses (measured via cardiac, pupil, and respiratory activity) during music-listening, only a small subset was in fact relevant to listeners’ on-line pleasure ratings. Despite that, we identified a weak mediation effect of physiology on shaping musical pleasure. Our results thus indicate that whilst musical pleasure may be embodied in bodily responses, the mapping between subjective experience and physiology is likely one-to-many—in line with psychological construction theories of emotion—and not one-to-one as is commonly assumed in classical basic emotion theories.

## Introduction

People in the industrialised world nowadays spend considerable amount of their waking hours listening to music (1, 2), primarily due to its ability to evoke strong emotions (3). While emotions are often accompanied by changes in autonomic neural responses (4), the role of physiology in our affinity for music is still unclear. Insights on the connection between autonomic responses and musical pleasure are crucial not only towards understanding what emotions are (5), but also have practical implications for music therapy (6) and the development of physiology-based, client-tailored music recommendation systems (7).

Whilst neuroimaging (8–12), neurostimulation (13), and pharmacological (14) studies have convincingly demonstrated that musical pleasure is subserved by dopaminergic signals in the mesolimbic reward network, the link between autonomic responses and musical pleasure has thus far been inconclusive. Although many studies have identified a relationship between listeners’ pleasure and their physiological responses derived from cardiac (15–27), respiratory (19, 21, 23), electrodermal (6, 13, 15, 20–23, 27, 28), pupil (29, 30), or facial muscle activity (6, 26, 27), the magnitude and direction of effects were often inconsistent. For example, while studies have generally reported a positive relationship between musical pleasure and heart rate (15, 16, 19–26), some studies detected no significant changes (6, 20) or a negative relationship (17, 18, 27). Likewise, skin conductance has been reported to be positively (6, 13, 15, 20–23, 28), negatively (27), and not significantly (19, 26, 28) related to musical pleasure.

One possibility for this discrepancy is that many of these studies used a combination of self- and experimenter-selected music as stimuli to investigate music-evoked frisson (15, 16, 20, 21, 23). Frisson is a psychophysiological response characterised by an intensely pleasurable chilling sensation that occurs at moments of peak emotional arousal (see (31) for a review), and studies used self-selected music to ensure that frisson were reliably evoked by subjects in the experiment. However, using both self- and experimenter-selected music could mean that the observed effects were confounded by familiarity. Indeed, music familiarity has also been shown to be related to changes in heart rate (17), skin conductance responses (32), and pupil activity (33), and could engage additional extra-musical processes such as episodic memory and visual imagery (34, 35).

Furthermore, very different musical stimuli were often used to evoke pleasure in previous studies. In addition to those using self- and experimenter-selected music, (18) used music from the Renaissance period and heavy metal as pleasant and unpleasant stimuli, respectively, (24) used tonal versus atonal music, whilst (29) chose music from multiple music genres as pleasure-evoking stimuli. Not only could the observed changes in autonomic responses have been due to large differences in acoustic properties within these diverse stimuli, listeners’ perception could also have been modified as they employ stylistic-dependent listening strategies. This is supported by studies showing changes in physiology in response to variations in tempo (36, 37) and consonance (38), effects of pitch on pleasure and arousal (39), as well as changes in harmonic expectancy when listening to rock and classical music (40).

To resolve these issues, we developed time-warped representational similarity analysis (twRSA) as a method to relate temporal changes in variable-duration stimuli across features derived from multiple modalities in terms of their representational structure. This enabled us to connect acoustic information in the musical stimuli to variations in listeners’ subjective pleasure and physiology whilst presenting subjects with only one composition but interpreted by multiple performers (thus with different durations). This alternative approach in stimulus selection allowed us to better control for stimulus confounds compared to the diverse musical excerpts used in previous studies. Using structural equation modelling, we further demonstrated that listeners’ autonomic responses mediated acoustic-evoked changes in musical pleasure.

## Results

### Generating acoustic, physiological, and behavioural representations from dynamic features

We recorded cardiac, respiratory, and pupil activity from 30 conservatory-level pianists as they listened to short musical excerpts and rated their subjective pleasure continuously using a mechanical slider. Unlike existing studies, our musical stimuli were all excerpted from the same composition (Variation XVIII from *Rhapsody of a Theme by Paganini*, Op. 43 by Sergei Rachmaninov) and comprised seven human and one MIDI interpretations (see Table 1). We also extracted 11 time-varying acoustic features that characterised the dynamical, rhythmic, spectral, and tonal properties of the eight musical expressions (refer to Table 3 for a description).

**Table 1:** Stimulus duration and reference information for the seven human performance recordings and one MIDI rendition of *Rhapsody of a Theme by Paganini*, Op. 43 by Sergei Rachmaninov from which the musical stimuli were derived. Note that only the solo piano was featured in the stimuli. Stimulus duration was trimmed to the nearest 0.001 s.

| *Soloist* | *Orchestra* | *Conductor* | *Duration* |
| --- | --- | --- | --- |
| Vladimir Ashkenazy | Philharmonia Orchestra | Bernard Haitink | 16.333 s |
| Van Cliburn | The Philadelphia Orchestra | Eugene Ormandy | 13.292 s |
| Andrei Gavrilov | The Philadelphia Orchestra | Riccardo Muti | 11.513 s |
| Zoltan Kocsis | San Francisco Symphony Orchestra | Edo de Waart | 11.270 s |
| Arthur Rubinstein | Chicago Symphony Orchestra | Fritz Reiner | 14.752 s |
| Arcadi Volodos | Rotterdam Philharmonic Orchestra | Valery Gergiev | 14.876 s |
| Lilya Zilberstein | RAI National Symphony Orchestra | Eliahu Inbal | 15.486 s |
| (MIDI) |  |  | 11.828 s |

We next sought to capture the representational geometry of temporal variations in each feature based on their (dis)similarity between the musical expressions. As direct computation was not possible since each expression differed in duration, we constructed warping paths that optimally aligned each expression pair with respect to power represented on the Mel-spectrogram (see Figure 1). Each warping path temporally transforms and matches signals recorded during presentation of the different expressions by their corresponding auditory content in a non-linear manner. This enabled us to compute the pairwise dissimilarity across each expression and to derive time-warped representational dissimilarity matrices (twRDMs) for each feature (see Figure 2).

**Figure 1:**
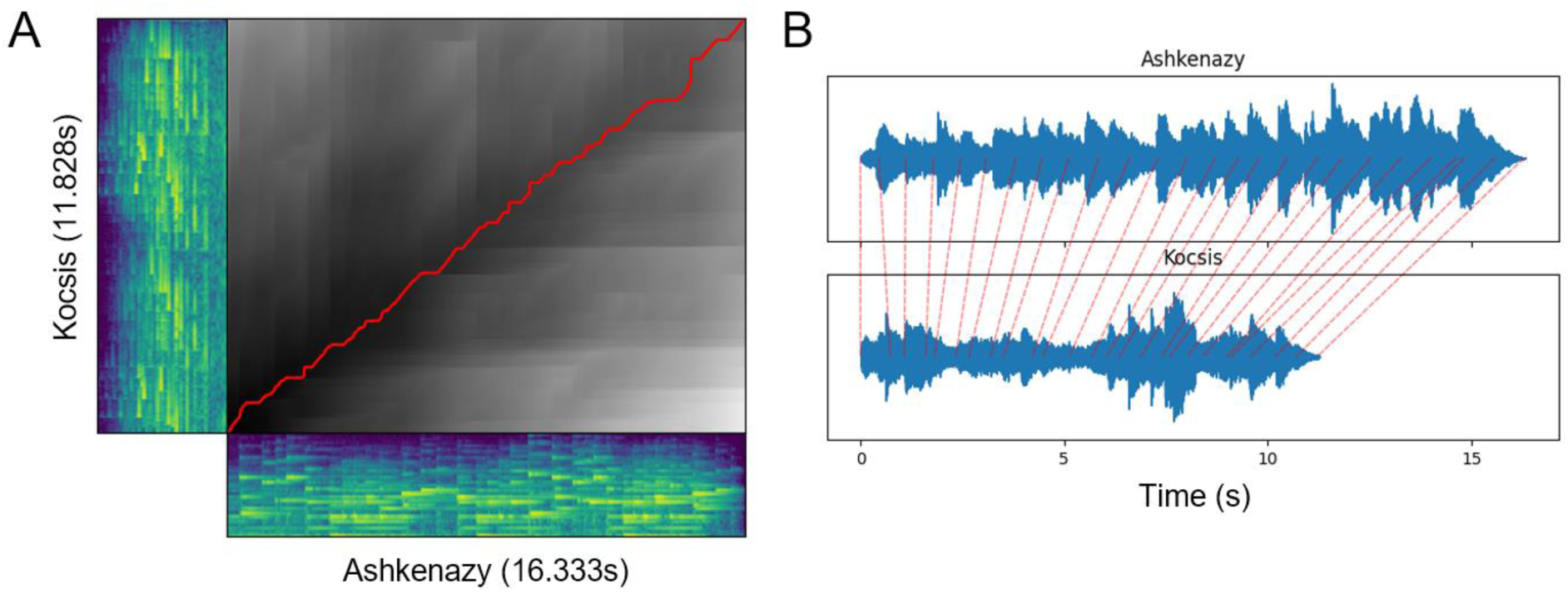
Mapping two stimuli with different durations using dynamic time warping (see Materials and Methods). (A) Colour shaded regions show the spectrogram of the longest and shortest musical excerpt (performed by Ashkenazy and Kocsis, respectively). Black shaded region shows the accumulated cost matrix in aligning the two expressions with respect to the cosine distance. Red line shows the optimal warping path with the minimum total cost. (B) Alternative wave-form representation of the two musical expressions. Dotted red lines denote the corresponding map between temporal locations of the two stimuli using the optimal warping path.

**Figure 2:**
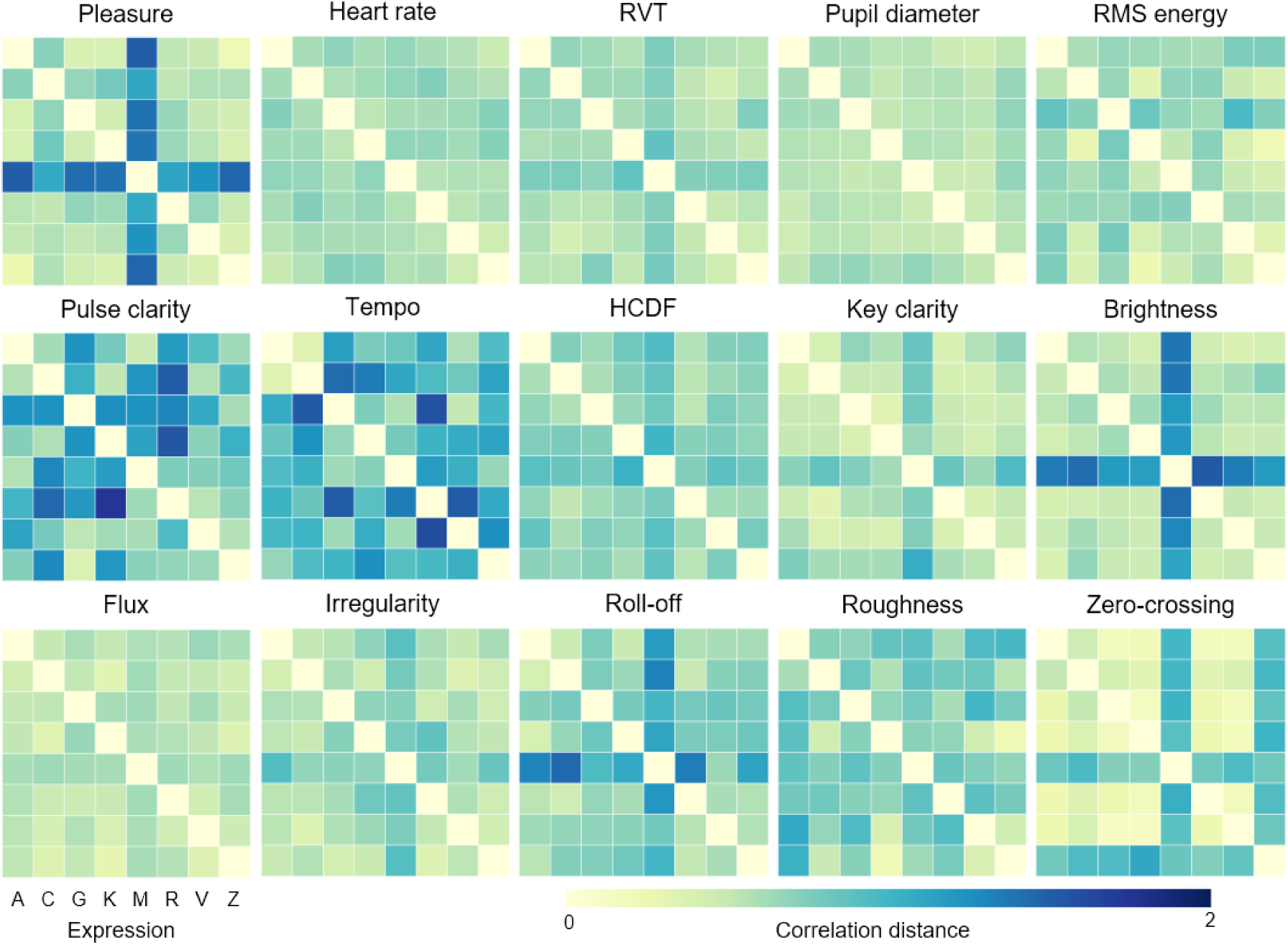
Time-warped representational dissimilarity matrices (twRDMs) for acoustic features and subject-averaged behavioural and physiological responses. Darker colours indicate higher correlation distance between expression pairs. Note that each twRDM is not symmetric unlike classical RSA as the optimal warping path between two expressions is asymmetric. Axis labels indicate the eight (7 human + MIDI) interpretations of the stimuli: A: Ashkenazy; C: Cliburn; G: Gavrilov; K: Kocsis; M: MIDI; R: Rubinstein; V: Volodos; Z: Zilberstein. RVT: respiration volume per time; HCDF: harmonic change detection function; RMS: root-mean-square.

### Variations in acoustic features were relevant to physiology, but only a subset was relevant to subjective pleasure

We first performed multidimensional scaling (MDS) to obtain a two-dimensional projection of the relative distance between each twRDM on the subject-averaged level (see Figure 3). This revealed that subject-averaged cardiac, respiratory, and pupil activity were closely clustered together with tonal and spectral acoustic features of the stimuli such as HCDF (harmonic change detection function), flux, and irregularity, whilst subject-averaged pleasure ratings were more related to brightness.

**Figure 3:**
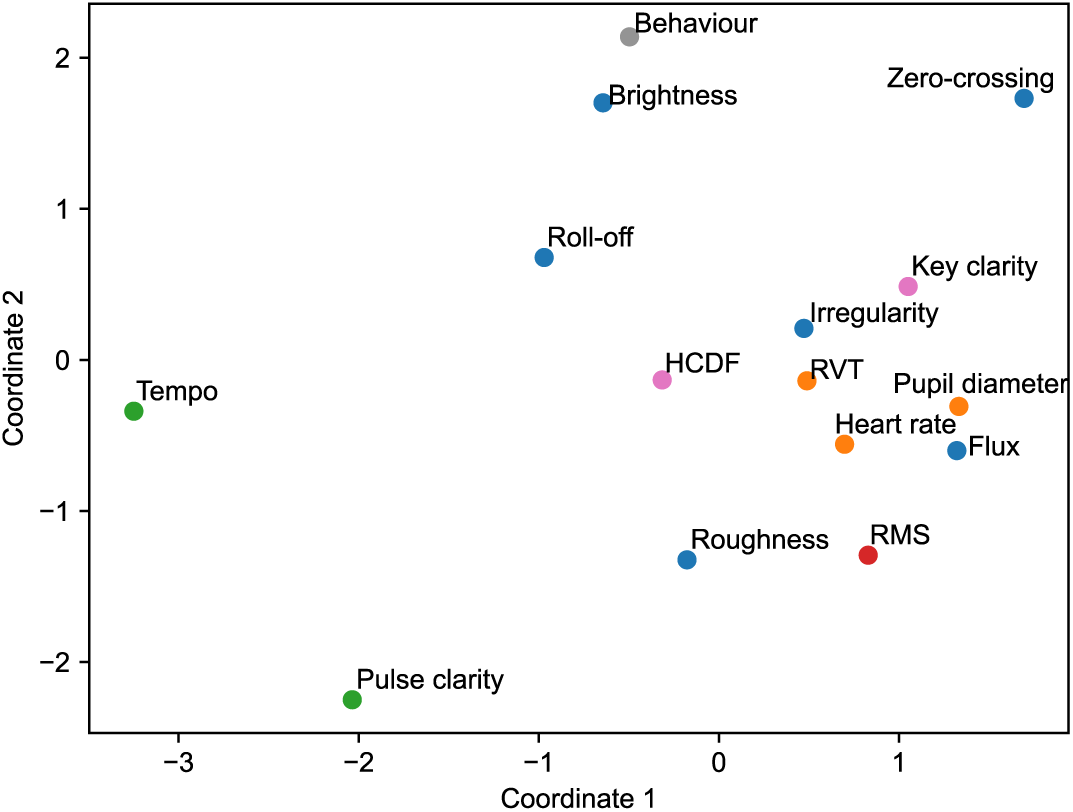
Multidimensional scaling plot showing relative representational similarity across acoustic features and subject-averaged responses in two dimensions. While multiple tonal and spectral features were similar to physiological responses, pleasure ratings were more related to brightness.

We next formally quantified the similarity between twRDMs representing temporal variations in each acoustic feature and subjects’ behavioural and physiological responses during music presentation using Pearson’s correlation (see Figure 4 and Table 2). Akin to results from multidimensional scaling, amongst the 11 acoustic features, HCDF, flux, and irregularity consistently showed the highest similarity with subjects’ cardiac, respiratory, and pupil activity. Brightness and roll-off were likewise most similar to subjects’ on-line pleasure ratings, whilst similarity of HCDF, flux, and irregularity were relegated to ranks 5, 6, and 3, respectively.

**Figure 4:**
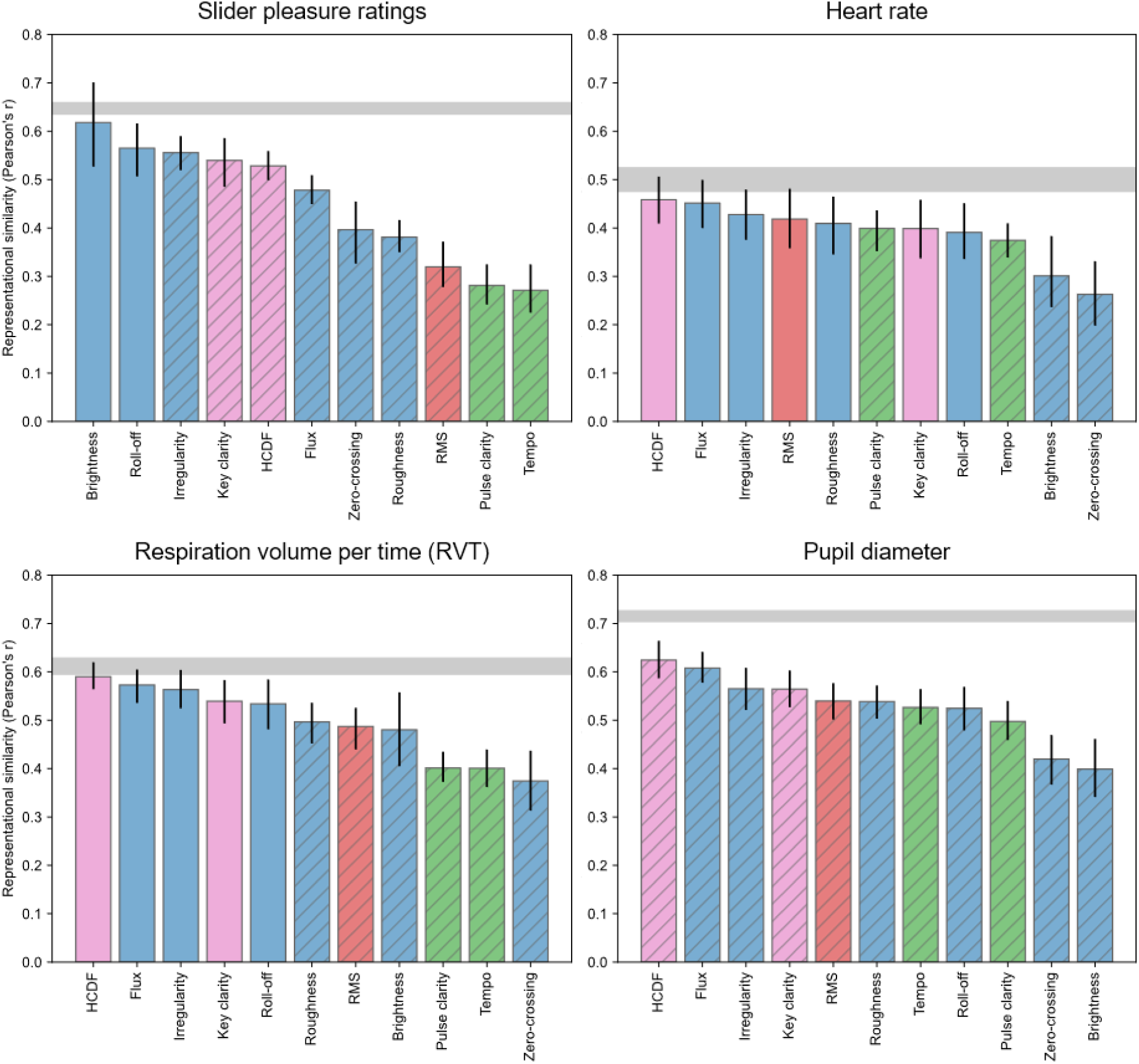
Comparing representational similarity across acoustic features and behavioural and physiological responses. Whilst the many acoustic features showed high representational similarity to physiological responses, only brightness and roll-off were relevant to subjects’ slider pleasure ratings. Grey regions indicate noise ceiling of behavioural and physiological response similarities. Dynamical, rhythmic, spectral, and tonal acoustic features are shown as red, green, blue, and pink bars, respectively. Solid bars indicate no significant difference between an acoustic feature and lower noise ceiling (based on one-sided Wilcoxon’s test after Bonferroni correction). Hatched bars denote significant differences. Error bars indicate 95% confidence intervals derived from 5,000 bootstrap samples.

**Table 2:**
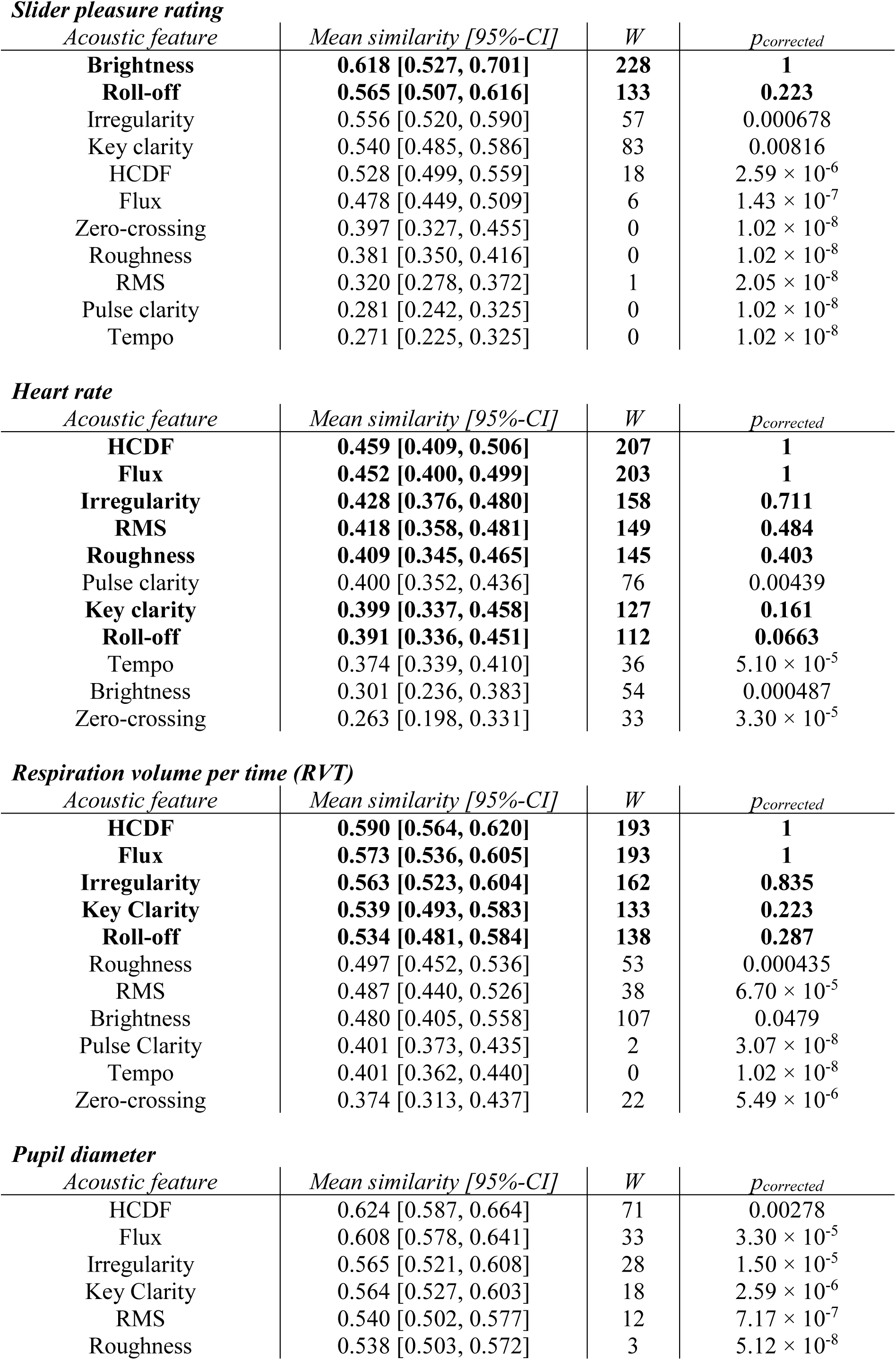
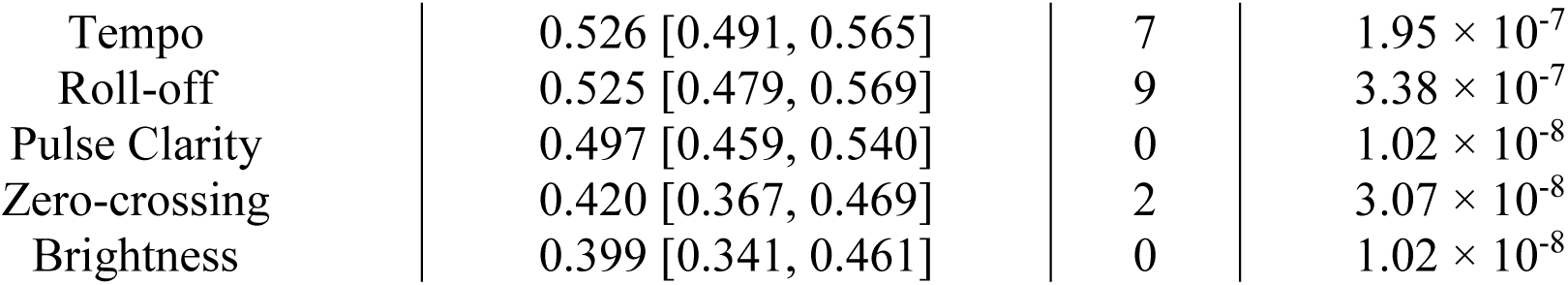
Representational similarity between changes in acoustic features and changes in behavioural and physiological responses, quantified using Pearson’s correlation. Acoustic features whose similarity did not differ significantly from the lower noise ceiling of each response modality (based on one-sample Wilcoxon’s tests after Bonferroni correction) are shown in **bold**. HCDF: harmonic change detection function; RMS: root-mean-square; W: test statistic.

| *Acoustic feature* | *Mean similarity [95%-CI]* | *W* | *p<sub>corrected</sub>* |
| --- | --- | --- | --- |
| **Brightness** | **0.618 [0.527, 0.701]** | **228** | **1** |
| **Roll-off** | **0.565 [0.507, 0.616]** | **133** | **0.223** |
| Irregularity | 0.556 [0.520, 0.590] | 57 | 0.000678 |
| Key clarity | 0.540 [0.485, 0.586] | 83 | 0.00816 |
| HCDF | 0.528 [0.499, 0.559] | 18 | $2.59 \times 10^{-6}$ |
| Flux | 0.478 [0.449, 0.509] | 6 | $1.43 \times 10^{-7}$ |
| Zero-crossing | 0.397 [0.327, 0.455] | 0 | $1.02 \times 10^{-8}$ |
| Roughness | 0.381 [0.350, 0.416] | 0 | $1.02 \times 10^{-8}$ |
| RMS | 0.320 [0.278, 0.372] | 1 | $2.05 \times 10^{-8}$ |
| Pulse clarity | 0.281 [0.242, 0.325] | 0 | $1.02 \times 10^{-8}$ |
| Tempo | 0.271 [0.225, 0.325] | 0 | $1.02 \times 10^{-8}$ |

*Heart rate*
| *Acoustic feature* | *Mean similarity [95%-CI]* | *W* | *p<sub>corrected</sub>* |
| --- | --- | --- | --- |
| **HCDF** | **0.459 [0.409, 0.506]** | **207** | **1** |
| **Flux** | **0.452 [0.400, 0.499]** | **203** | **1** |
| **Irregularity** | **0.428 [0.376, 0.480]** | **158** | **0.711** |
| **RMS** | **0.418 [0.358, 0.481]** | **149** | **0.484** |
| **Roughness** | **0.409 [0.345, 0.465]** | **145** | **0.403** |
| Pulse clarity | 0.400 [0.352, 0.436] | 76 | 0.00439 |
| **Key clarity** | **0.399 [0.337, 0.458]** | **127** | **0.161** |
| **Roll-off** | **0.391 [0.336, 0.451]** | **112** | **0.0663** |
| Tempo | 0.374 [0.339, 0.410] | 36 | $5.10 \times 10^{-5}$ |
| Brightness | 0.301 [0.236, 0.383] | 54 | 0.000487 |
| Zero-crossing | 0.263 [0.198, 0.331] | 33 | $3.30 \times 10^{-5}$ |

*Respiration volume per time (RVT)*
| *Acoustic feature* | *Mean similarity [95%-CI]* | *W* | *p<sub>corrected</sub>* |
| --- | --- | --- | --- |
| **HCDF** | **0.590 [0.564, 0.620]** | **193** | **1** |
| **Flux** | **0.573 [0.536, 0.605]** | **193** | **1** |
| **Irregularity** | **0.563 [0.523, 0.604]** | **162** | **0.835** |
| **Key Clarity** | **0.539 [0.493, 0.583]** | **133** | **0.223** |
| **Roll-off** | **0.534 [0.481, 0.584]** | **138** | **0.287** |
| Roughness | 0.497 [0.452, 0.536] | 53 | 0.000435 |
| RMS | 0.487 [0.440, 0.526] | 38 | $6.70 \times 10^{-5}$ |
| Brightness | 0.480 [0.405, 0.558] | 107 | 0.0479 |
| Pulse Clarity | 0.401 [0.373, 0.435] | 2 | $3.07 \times 10^{-8}$ |
| Tempo | 0.401 [0.362, 0.440] | 0 | $1.02 \times 10^{-8}$ |
| Zero-crossing | 0.374 [0.313, 0.437] | 22 | $5.49 \times 10^{-6}$ |

Table 2: Representational similarity between changes in acoustic features and changes in behavioural and physiological responses, quantified using Pearson’s correlation.
| *Acoustic feature* | *Mean similarity [95%-CI]* | *W* | *p<sub>corrected</sub>* |
| --- | --- | --- | --- |
| HCDF | 0.624 [0.587, 0.664] | 71 | 0.00278 |
| Flux | 0.608 [0.578, 0.641] | 33 | $3.30 \times 10^{-5}$ |
| Irregularity | 0.565 [0.521, 0.608] | 28 | $1.50 \times 10^{-5}$ |
| Key Clarity | 0.564 [0.527, 0.603] | 18 | $2.59 \times 10^{-6}$ |
| RMS | 0.540 [0.502, 0.577] | 12 | $7.17 \times 10^{-7}$ |
| Roughness | 0.538 [0.503, 0.572] | 3 | $5.12 \times 10^{-8}$ |
| Tempo | 0.526 [0.491, 0.565] | 7 | $1.95 \times 10^{-7}$ |
| Roll-off | 0.525 [0.479, 0.569] | 9 | $3.38 \times 10^{-7}$ |
| Pulse Clarity | 0.497 [0.459, 0.540] | 0 | $1.02 \times 10^{-8}$ |
| Zero-crossing | 0.420 [0.367, 0.469] | 2 | $3.07 \times 10^{-8}$ |
| Brightness | 0.399 [0.341, 0.461] | 0 | $1.02 \times 10^{-8}$ |

Next, we estimated the noise ceiling of subjects’ behavioural and physiological twRDMs to derive an upper bound on the maximum representational similarity with subjects’ responses given the variability in the data. This is the correlation that would have been detected for a hypothetical feature that generated the observed data. We identified several tonal and spectral acoustic features whose similarity did not differ statistically from the lower noise ceilings of cardiac and respiratory activity based on one-sided Wilcoxon’s test after Bonferroni correction, namely HCDF, flux, irregularity, key clarity, and roll-off (see Table 2). In addition, root-mean-square (RMS) energy and roughness did not differ from the cardiac noise ceiling, whilst representational similarity of all acoustic features differed significantly from the pupil noise ceiling. However, only brightness and roll-off did not differ significantly from the behavioural noise ceiling. These suggest that though many acoustic features could explain subjects’ physiological responses during music listening, only a few were relevant to their subjective pleasure.

### Physiological responses weakly shape spectral-driven variations in musical pleasure

Whilst comparisons with the noise ceiling revealed that variations in spectral features were separately related to changes in behaviour and physiology, it was not clear whether subjects’ time-varying pleasure ratings could have been shaped by their physiological responses, or vice versa. To this end, we used multilevel structural equation modelling to test the effect of physiology on mediating the changes in musical pleasure as driven by spectral variations in the musical stimuli. One advantage of structural equation modelling over standard regression techniques is the ability to examine effects of latent variables that are derived from observed variables, rather than observed variables themselves. We were therefore able to model ‘physiology’ as one latent variable composed of subjects’ cardiac, respiratory, and pupil twRDMs, and ‘acoustics’ as another composed of brightness and roll-off twRDMs, and ‘pleasure’ as another based on subjects’ behavioural twRDMs.

As shown in Figure 5, this analysis revealed a significant direct effect of acoustics on shaping musical pleasure (γ_standardised_ = 0.486, γ_unstandardised_ = 0.604, 95%-bootstrapped CI = [0.409, 0.683], p < 0.0005) and physiology (γ_standardised_ = 0.716, γ_unstandardised_ = 1.026, 95%CI = [0.932, 1.337], p < 0.0005), as well as physiology on pleasure (β_standardised_ = 0.142, β_unstandardised_ = 0.123, 95% CI = [0.082, 0.254], p < 0.0005). We additionally observed a significant indirect effect of acoustics on musical pleasure via physiology (γ_standardised_ = 0.101, γ_unstandardised_ = 0.126, 95%CI = [0.092, 0.291], p < 0.0005), thus yielding a total standardised effect of 0.587 (γ_unstandardised_ = 0.731, 95%CI = [0.666, 0.810], p < 0.0005) on pleasure. Examining the physiological latent variable further revealed a predominant contribution of respiratory activity (λ_standardised_ = 0.708) compared to pupil and cardiac activity (λ_standardised_ = 0.531 and 0.458, respectively), whilst both brightness and roll-off received similar loadings for the acoustic latent variable (λ_standardised_ = 0.943 and 0.950, respectively). While these findings support the role of physiology in shaping musical pleasure, it should be noted that the mediation effect was small, explaining only ∼17% variance.

**Figure 5:**
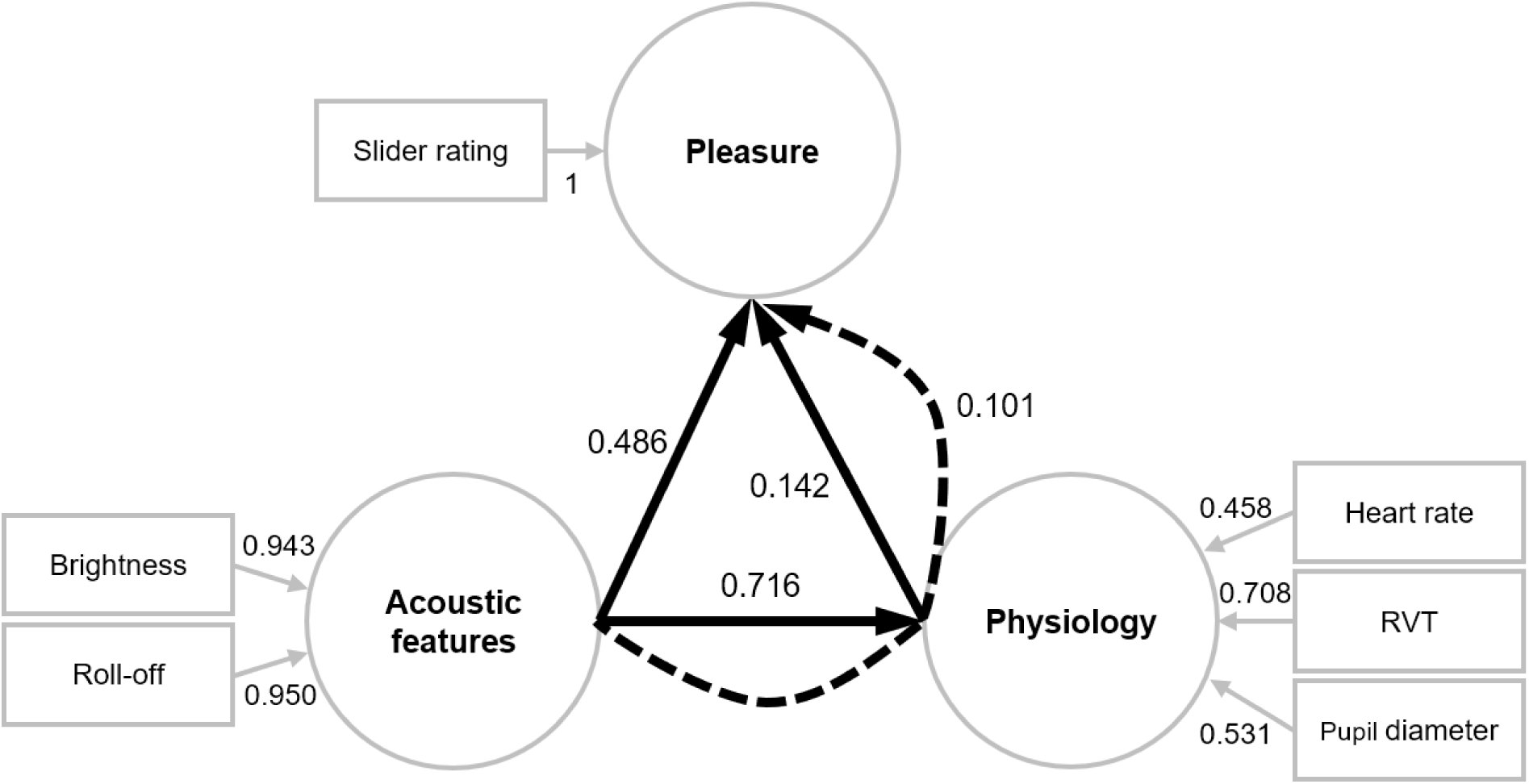
Structural equation modelling revealed that the effect of spectral acoustic features on modulating musical pleasure was mediated by subjects’ physiological responses. Circles denote latent variables; boxes denote observed variables. Numbers next to solid black arrows indicate size of standardised direct effects between latent variables. Number next to dotted arrow denotes size of standardised indirect effect of acoustics on musical pleasure via the mediation of physiology. Numbers next to solid grey arrows indicate standardised loadings of latent variables by each observed variable.

### Control analyses

Given the distinct difference in similarity between human expressions and MIDI (which had no rhythm or loudness variations), we repeated the above analyses with MIDI removed. First, we observed that in addition to brightness and roll-off, HCDF and irregularity also did not differ significantly from the behavioural lower noise ceiling (see Supplementary Figure 1). This implies that changes in HCDF and irregularity could also now viably predict changes in behaviour. Second, while the similarity between acoustic features and physiology were comparable to those with MIDI included, brightness additionally did not differ significantly from the pupil and cardiac lower noise ceilings. Third, in our structural equation modelling analysis with MIDI excluded (see Supplementary Figure 2), we observed a larger direct effect of physiology on musical pleasure (0.342 vs 0.142), as well as a larger indirect effect of acoustics on pleasure via physiology (0.298 vs 0.101). Taken together, these provide strong support that our results were not driven by the MIDI expression, and that its inclusion likely resulted in more conservative findings.

Furthermore, respiratory sinus arrhythmia is a phenomenon commonly observed in vertebrates in which heart rate fluctuates synchronously with each respiratory cycle (41). To rule out the possibility that the heart rate and RVT timeseries used to derive the twRDMs were confounded by respiratory sinus arrhythmia, we tested for the temporal relationship between the two modalities in each trial for all subjects. As shown in Supplementary Figure 3, we only identified a small portion of trials (34.2%) in which RVT in earlier time lags could explain subsequent heart rate (and not vice versa), with no significant difference across the eight expressions (χ^2^(7) = 10.4, p = 0.166). By contrast, directionality was inconclusive in 40.5% of trials, or reversed in 24.8% of trials. These suggest a minimal influence of respiration on cardiac activity in the current data.

## Discussion

Music evokes subjective emotional and physiological responses from listeners. Here, we introduced time-warped representational analysis (twRSA) as a method to relate acoustic changes in variable-duration stimuli to dynamic changes in subjective pleasure and autonomic neural responses during music-listening. We identified several tonal, spectral, and dynamical acoustic features, including HCDF (harmonic change detection function), key clarity, brightness, roll-off, and root-mean-square energy as plausible candidates in shaping subjects’ physiological responses. However, only two spectral features—brightness and roll-off—were relevant to subjects’ on-line pleasure ratings. Structural equation modelling further revealed a weak mediation of physiology on the effect of acoustic variations in shaping pleasure. While these results support an embodied view of musical pleasure, it suggests that the mapping between physiology and pleasure is likely not unique.

The dissociation between physiology and pleasure suggested from our findings support psychological construction theories (PCT) of emotion (51–53). Contrary to classical basic emotion theories (BET) (54–58) that assume a prototypical expression and neural signature corresponding to each emotional category (53, 55), PCT posit that emotions are instantiated concepts that arise as a response to predictions to the sensory environment based on previous idiosyncratic experiences (52). PCT further argue for a degeneracy account, where different neural processes—depending on context—interact and integrate to give rise to the same emotional percept and expression (51, 59). In line with this, akin to pleasant music, unpleasant music was likewise shown to evoke increased heart rate, and skin conductance responses compared to neutral music (27). Furthermore, a cruder, dimensional representation of emotion in terms of valence and arousal could better characterise listeners’ perceived emotions elicited by music compared to discrete emotional categories (68). Taken together, these corroborate the view that the correspondence between musical pleasure and evoked autonomic responses is likely not one-to-one, contrary to what is crucially assumed by many physiology-based music-evoked emotion classifiers (60–67).

One important corollary of our finding is that physiological and subjective pleasure responses to music might rely on distinct mechanisms. This view is supported by recent pharmacological evidence, where administration of oxycodone (an opioid agonist) and naltrexone (an opioid antagonist) resulted in respective increases and decreases in skin conductance responses to pleasant music, but no significant changes in subjective ratings (69). Similarly, another study (70) found that administration of naltrexone resulted in lower pleasure ratings but no significant changes in heart rate and respiration rate.

While our results speak for separate mechanisms between music-evoked pleasure and autonomic responses, they nevertheless also highlight their interaction. This is demonstrated by our structural equation modelling analyses, which showed that subjects’ physiological responses mediated the effects of acoustics on perceived pleasure. This finding supports embodied or enactivist (71) views of music-evoked emotions, which argue that emotional percepts are informed by bodily sensations (72, 73), and reinforce the proposal that music and the body are intimately connected (74). Interestingly, recent evidence using body sensation mapping have shown that different chord progressions (75) and acoustic features in music elicited emotions and bodily sensations that were consistent across cultures (72). This view is moreover supported by neuroimaging studies showing the encoding of music-evoked emotions in somatosensory (46) and motor cortices (76, 77), observed spontaneous movements such as ‘tapping to the beat’ in response to musical stimuli (78, 79) that is already manifested from 5 months of age (80), as well as embodied experiences to aversive music (81).

A key difference between the current study and previous work is that instead of merely examining the correlation between music-evoked pleasure and autonomic neural responses, here we identified plausible acoustic features within the musical stimuli that modulated both modalities. With this triage, we found a dissociation between musical acoustic features that predicted changes in physiological responses (i.e., HCDF, key clarity, brightness, roll-off, and root-mean-square energy), and those that predicted changes in subjective pleasure ratings (brightness and roll-off). In line with this, HCDF, key clarity and root-mean-square energy have been previously identified as acoustic factors relevant for classifying between music-evoked chills and tears and emotionally neutral episodes up to 8s before onset (42). On the other hand, roll-off has been associated with being joyfully moved by music (43). Likewise, brightness has been shown to be correlated with activity in the superior temporal gyrus and Heschl’s gyrus in the auditory cortex (44, 45), which are thought to generate acoustic-evoked emotional percepts (46) and whose activity (10, 47) and functional connectivity (8, 11) with the mesolimbic reward network were shown to be related to musical pleasure.

Another distinction is our focus on the relationship between acoustics, pleasure, and physiology *over time*. This circumvents the need to define an appropriate time-lag between two modalities *a priori*. Rather than relating acoustic features to pleasure or physiological responses at discrete time points, here we examined to what extent the temporal variations in acoustic features were similar to temporal variations in subjects’ pleasure ratings and physiological responses in terms of their representational (dis)similarity matrices. Our findings thus did not imply that stimuli with a higher brightness or roll-off were more preferred, but that changes in brightness and roll-off were related to changes in subjects’ pleasure ratings. Capturing dynamic changes in pleasure is meaningful not only because music is a continuous stimulus (10, 48), but also that listeners’ engagement is also thought to vary dynamically throughout a piece (49, 50). This is demonstrated in our data, where despite a very high Pearson’s correlation between listeners’ mean slider rating and their subsequent overall rating (r = 0.796, 95%-CI = [0.783, 0.808], p < 1.00 × 10^-5^), we observed a much weaker direct effect between listeners’ physiological responses and overall pleasure ratings, as well as a much weaker mediation effect of physiology on pleasure (see Supplementary Figure 4).

An unexpected finding was that the representation similarity of all acoustic features was significantly below the pupil lower noise ceiling. This indicated that none were plausible candidates in explaining the change in subjects’ pupil size during music stimulation. One possibility could be due to the stimuli presented in the current experiment. Unlike the diverse sets of compositions used to evoke different levels of pleasure in previous studies, we controlled our stimuli by presenting subjects with the same musical piece but interpreted by multiple performers. Differences in artistic expression across our stimuli might have been too subtle to evoke significant changes in pupil activity, which is thought to primarily reflect arousal or attention via sympathetic neural activity of the locus coeruleus-noradrenaline system (82) and modulated by activity in the internal layer of the superior colliculus and pretectal olivary nucleus (83). In line with this, pupillary responses to music have been shown to correlate with listeners’ arousal and tension ratings (84), and with music-evoked pleasure via interactions with predictability (30). The lack of variation in evoked arousal in the stimuli could further explain why we did not observe significant changes in skin conductance responses across our stimuli in our pilot experiment.

There are a few limitations in the current study. First, while our mediation analysis showed that acoustical features modulated musical pleasure via the influence of autonomic responses, it is important to note that the causality implied was purely statistical and not biological. While our attempt at ‘reverse mediation testing’ (i.e., interchanging the mediator and predictor, see Supplementary Figure 5) suggested a more plausible mediation effect of physiology on pleasure compared to pleasure on physiology, this approach has been argued to lead to spurious conclusions (85, 86). Instead, future experiments could adopt a *moderation-of-process* design (87), where physiology (e.g., breathing rate) is explicitly manipulated during music-listening via biofeedback. This intervention would provide direct evidence on the causal role of physiology on musical pleasure. Second, we recruited conservatory-level pianists as subjects since we posited that a certain degree of expertise was required to discern the nuanced differences in our stimuli. In terms of the BRECVEMA framework of mechanisms underlying music-evoked emotions (88), we speculate that our stimuli evoked pleasure primarily through *aesthetic judgment* and *contagion*. By contrast, music that activated *brain stem reflex* and *episodic memory* mechanisms—which generally do not require musical expertise—have been shown to robustly elicit skin conductance responses and changes in facial muscle activity (89, 90). To what extent each mechanism engages autonomic neural activity during music-listening remains to be shown. Lastly, although we examined the relationship between evoked pleasure and physiological responses with respect to variations in acoustic features in the musical stimuli, listeners’ preferences are also thought to be shaped by extra-musical factors such as personality and culture (91). Future work could explore whether such contextual modifiers serve as a common denominator that explain music-evoked pleasure and its accompanying autonomic responses.

## Materials and Methods

### Subjects

Data were collected from 30 healthy adults (26 females, age: M = 22.4 y, SD = 2.55 y) with conservatory-level training in piano performance (M = 18.2 y) and who were still actively playing the instrument at the time of the experiment. Subjects reported normal hearing, no known history of neuropsychological disorders, hypertension, or cardiovascular disease. Subjects who normally wore glasses either wore contact lenses or took them off for the experiment. Sample size was chosen based on previous studies showing physiological differences in music preference (6, 13, 15–18, 20–22, 24, 26–30, 38, 92, 93), and the sampling strategy was based on subject availability.

The study was approved by the Sony Bioethics Committee and all subjects gave their written informed consent prior to the experiment. This study was not preregistered.

### Stimuli

Musical stimuli consisted of seven human performance recordings (refer to Table 1 for reference information) and a MIDI rendition of a short piano solo excerpted from Variation XVIII: Andante cantabile (mm. 639–644) from Sergei Rachmaninov’s *Rhapsody of a Theme by Paganini*, Op. 43 (duration M = 13.668 s, SD = 1.962 s). The MIDI stimulus had neither loudness nor rhythm deviations across notes. The eight stimuli were sliced (with 1 s fadeout introduced) and normalised in volume using Audacity 3.13 (www.audacityteam.org), and were played back from wave files in stereo at 16 bits per sample and a sampling rate of 44.1 kHz.

### Physiological data collection

Cardiac and respiratory signals were recorded using a Polymate Mini A108 (Miyuki Giken Co., Ltd., Tokyo, Japan) wireless physiological measurement device. To record cardiac activity, we used a three-lead placement with Ag-Cl electrodes placed on both wrists and on the left ankle. To measure the extent of chest expansion during respiration, a respiratory belt transducer was attached around the sternum. Pupil diameter was recorded using a Pupil Core eye tracker (Pupil Labs GmbH, Berlin, Germany) mounted over subjects’ nose. To ensure accurate estimation, eye cameras were adjusted such that each pupil was placed as close to the central field of view as possible and calibrated before the start of the experiment.

All physiological signals were recorded at a sampling rate of 1 kHz.

### Procedure

Subjects were seated comfortably in the experiment room in front of a 17-inch LCD monitor with the physiological sensors attached.

Each trial began with a fixation cross displayed in the centre of the screen for which subjects were instructed to focus on during the trial. One of the eight musical expressions was presented 1 s afterwards at a comfortable volume using in-ear earphones. Upon stimulus onset, subjects continuously rated how much they liked the musical excerpt using a custom-built mechanical slider held in their right hand (sampling rate = 1 kHz). The slider had notches on each integer within the interval [1, 7] and subjects were asked to adjust the knob of the slider with their right thumb to the corresponding pleasure rating throughout the excerpt, with the rating ‘1’ representing ‘disliking very much’ and ‘7’ representing ‘liking very much’. After stimulus presentation, subjects gave an overall pleasure rating of the excerpt on a [-50,50] integer-scale using the mouse, with ‘-50’ indicating ‘very much disliked’ and ‘50’ indicating ‘very much liked’. Next, the slider was reset to ‘4’ and the following trial ensued.

Each block consisted of eight trials, with each expression presented once in a randomised order, and a total of 15 blocks were presented. After each stimulus presentation in the first block, subjects also provided binary feedback (Yes/No) on whether they were familiar with each excerpt, to which all responded negative.

### Data preprocessing

Cardiac and respiratory data were preprocessed using the PhysIO toolbox (94) for noise removal and feature extraction. Cardiac data were first bandpass-filtered using a Chebyshev Type 2 filter between 0.3 and 9 Hz. Then R-R intervals between two successive cardiac cycles were extracted using robust peak detection and converted to instantaneous heart rate (HR) by taking the reciprocal from a 6 s-moving average window.

Respiration data underwent outlier rejection at a threshold of 5 SD (due to asymmetric distributions) and despiking using a 0.5s sliding-window median filter. This was followed by band-pass filtering with an order 20-infinte impulse response (IIR) filter between 0.01 and 2 Hz, and normalisation to [-1, 1]. Next, differences in maxima and minima in the normalised respiration data were used to obtain the volume of each respiratory cycle, which was then divided by the breath duration (time between two successive maxima) to obtain respiration volume per time (RVT). RVT timeseries have the advantage of capturing changes in both depth and rate information during respiration and are also highly correlated with end-tidal CO_2_ (95), and was therefore used to measure respiratory activity in the current study.

The resulting HR and RVT timeseries were then epoched to each stimulus presentation time-window for every trial, then transformed into z-scores within each trial, and set to 0 at stimulus onset using a subtractive baseline.

Pupil diameter was first extracted from a 3D eye model estimated from the pupil camera using the eye tracker’s *Pupil Player* software, with eyeblinks and motion artefacts rejected at a confidence level of 0.5 and filter threshold of 200 ms. The data were then mean-averaged across both eyes and epoched to each stimulus presentation time-window for every trial, and were rejected if the portion of eyeblinks during stimulus presentation exceeded 0.5. Finally, following recommendations in (96), missing data from eyeblinks were imputed using cubic spline interpolation, and the pupil diameter timeseries were set to 0 at stimulus onset using a subtractive baseline correction after transformation to z-scores within each trial.

The epoched HR, RVT, and pupil timeseries, as well as slider ratings were then down-sampled to 100 Hz to conserve disk space.

### Acoustic feature extraction

We extracted 11 time-varying acoustic features present in the MIR toolbox (97) from the eight musical stimuli. These features (summarised in Table 3 below) were chosen to capture changes in dynamics, tonality, rhythm, and spectral properties within the stimuli. The features were then resampled to 100 Hz using linear interpolation.

**Table 3:**
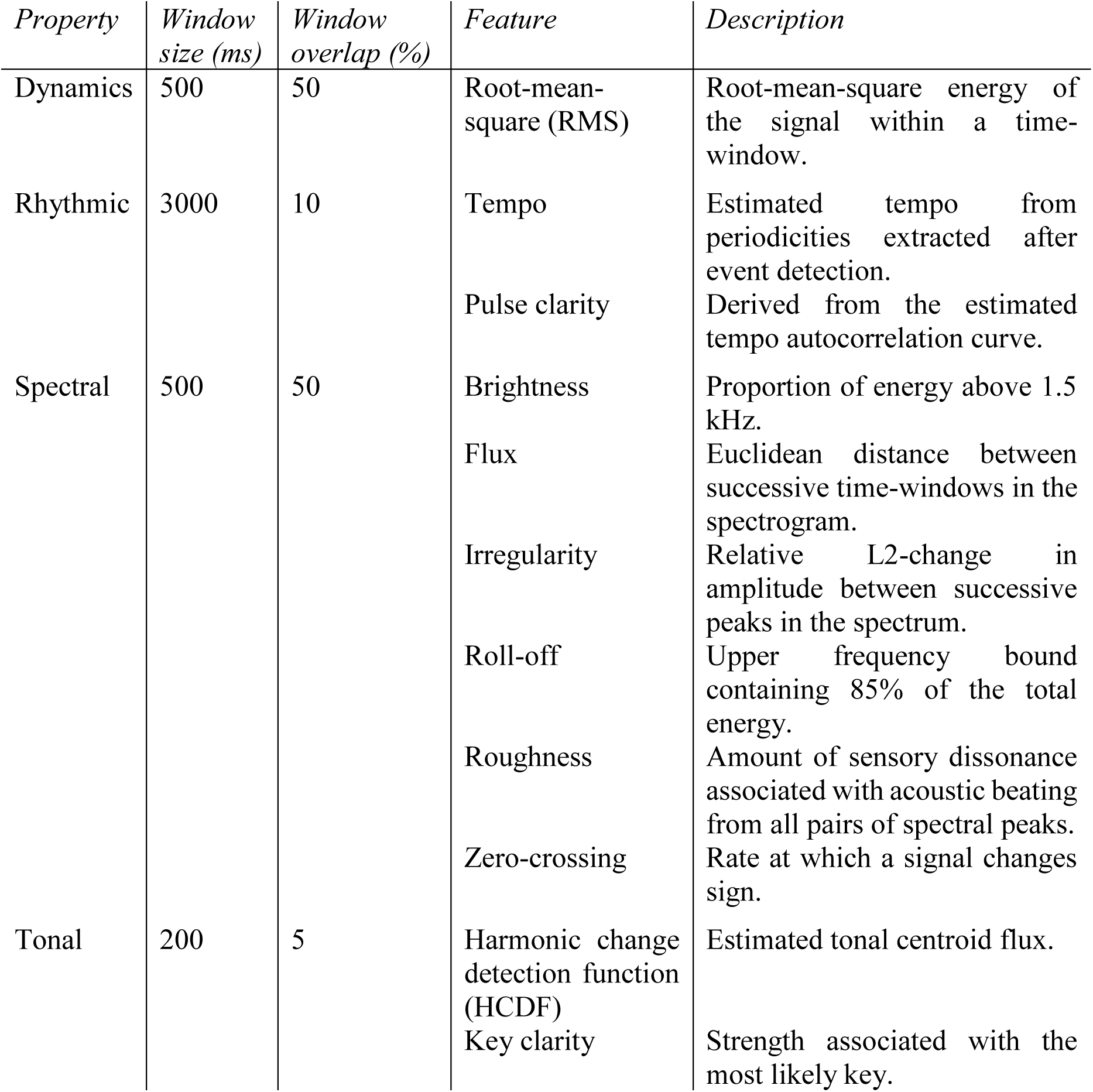
Summary description of time-varying acoustic features extracted from the stimuli.

### Time-warped representational similarity analysis (twRSA)

We developed time-warped representational similarity analysis (twRSA) to link temporal changes across multiple modalities in our variable-duration stimuli. twRSA is an extension of representational similarity analysis (RSA), which is a framework originally devised to quantify the correspondence between neural activity, behavioural measurements, and computational modelling (98–101). The key insight of RSA is leveraging similarity across each stimulus pair to characterise the representational geometry of a given modality, and then comparing these representations across modalities. While similarity across stimulus pairs is typically computed across spatial features in RSA (e.g., BOLD activity across voxels in a brain region, amplitude across EEG sensors at a specific timepoint, latent embeddings of specific layers in a neural network), it is possible to use time-varying features to compute pairwise similarity in RSA (for example, by taking the correlation between two timeseries (102)). However, a critical caveat to this approach has been that all stimuli must have identical duration (or more generally, the same number of features). That is because computing pairwise similarity requires an explicit map between each feature in the two stimuli.

To overcome the need for identical stimulus duration, one approach is to compute similarity based on summary statistics (such as the mean and variance). However, meaningful variations in the signal are likely lost as they are reduced into single values. A better approach, which we propose here, is to construct an explicit mapping between timepoints across stimuli using dynamic time warping.

### Dynamic Time Warping (DTW)

Let *X* = (*x*_1_, …, *x*_*N*_) and *Y* = (*y*_1_, …, *y*_*M*_) be two timeseries. The goal of DTW is to find an *optimal warping path P*\* where given some optimality condition, every timepoint in *X* is mapped to a corresponding timepoint in *Y* and vice versa. Below, we illustrate how to obtain *P*\* via dynamical programming. Please refer to (103) for the full derivation of the algorithm.

We begin by defining a non-negative function *c* to describe the dissimilarity, or *local cost*, between a timepoint in each timeseries within a feature space *F*. This allows us to compute the *N* × *M cost matrix C* such that *C*(*i*, *j*) = *c*(*x*_*i*_, *y*_*j*_).

Next, we define a map *P* = (*p*_1_, …, *p*_*L*_), *p*_*k*_ = (*n*_*k*_, *m*_*k*_) for *n*_*k*_ ≤ *N* and *m*_*k*_ ≤ *M* that satisfies the following constraints:

1. *p*_1_ = (1,1) and *p*_*L*_ = (*N*, *M*), and
2. *p*_(*i*+1)_ − *p*_*i*_ ∈ {(1,0), (0,1), (1,1)}.

*P* is referred to as a *warping path* and associates one timepoint in *X* with another in *Y* (and vice versa) with the condition that 1) the start and end of both timeseries are aligned, and 2) that the mapping is monotonic (i.e., it cannot go back in time) and smooth.

This allows us to compute the *total cost* of *P*, which is given by 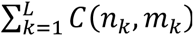, or the sum of local costs associated with traversing *P*. The optimal warping path *P*\* is then defined as the one with the minimum total cost.

Next, we construct the *N* × *M accumulated cost matrix D* such that 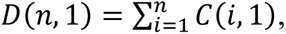 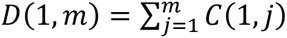, and *D*(*n, m*) = *C*(*n, m*)+*min*{*D*(*n*-1, *m*-1), *D*(*n*-1, *m*), *D*(*n*, *m*-1)} for *n*=2, …, *N* and *m* = 2, …, *M*.

Finally, we derive *P* ∗= (*p*_1_, …, *p*_*L*_) by the following algorithm:

1. Let *l* = 1 and *q*_1_ = (*N*, *M*)
2. Do *l* = *l* + 1 and *q*_*l*−1_ = (*n*, *m*) If *n* = 1 then *q*_*l*_ = (1, *m* − 1) Else if *m* = 1 then *q*_*l*_ = (*n* − 1,1) Else *q*_*l*_ = *argmin*_*n*,*m*_{*D*(*n* − 1, *m* − 1), *D*(*n* − 1, *m*), *D*(*n*, *m* − 1)} Until *q*_*l*_ = (1,1)
3. Then set *L* = *l* and return *P* ∗= (*q*_*L*_, …, *q*_1_) = (*p*_1_, …, *p*_*L*_).

In this study, DTW was carried for each stimulus pair using librosa 0.10.2 (104). The feature space *F* was defined to be the power on the Mel-spectrogram of each stimulus obtained using discrete Fourier transform (128 Mel bins, 2048 windows per sample, hop-length equal to 441, corresponding to 10 ms resolution). Upon inspection, we selected cosine similarity as the local cost function that best aligned two musical expressions, with the exception of Volodos-MIDI where Euclidean distance offered a superior alignment.

### Deriving time-warped representational dissimilarity matrices (twRDMs)

Having obtained the optimal warping path for each stimulus pair, the next step was to evaluate their temporal (dis)similarity after alignment. As in RSA, for each subject, we first mean-averaged their slider pleasure rating, HR, RVT, and pupil diameter timeseries across trials for each expression to improve signal-to-noise ratio. Next, for each modality, we obtained a time-warped representational dissimilarity matrix (twRDM) by computing the correlation distance between the aligned timeseries of each expression pair for every subject. Given timeseries *X* and *Y*, the correlation distance *d* is defined as

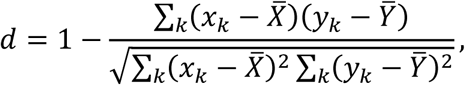

where 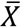 and 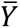 denote the sample mean of *X* and *Y*, respectively. A correlation distance of 0 suggests that two timeseries are perfectly positively linearly correlated, whilst the (maximum) distance of 2 suggests that they are perfectly negatively linearly correlated. However, note that unlike standard RSA, the resultant twRDM is not symmetric as the optimal warping path is non-linear.

### Testing similarity between twRDMs

To quantify the similarity between twRDMs, we computed Pearson’s correlation between the flattened twRDMs of each modality for all subjects. As each twRDM is non-symmetric, all entries in the matrix were used (unlike RSA, where only the upper/lower-triangle is used due to symmetry). Note that our interest was only in the size and not in the significance of these correlations, as their p-values are not meaningful due to the presence of temporal autocorrelations.

We furthermore computed the noise ceiling for each behavioural (i.e., slider pleasure rating) and physiological (i.e., HR, RVT, and pupil diameter) twRDM. The noise ceiling provides an upper bound in similarity based on an estimate of variability present across subjects (105). In particular, the upper noise ceiling is computed by taking the mean correlation between each subject’s z-transformed twRDM and the group-averaged twRDM after z-transformation, whilst the lower noise ceiling is defined as the mean correlation between each subject’s z-transformed twRDM and the group-averaged and z-transformed twRDM with that subject excluded (99).

### Structural equation modelling

Structural equation modelling (SEM) is a linear modelling framework that allows one to test statistically causal relationships between latent variables via regression (106). An important feature of SEM is the distinction between measurement models and structural models. Measurement models describe the connection between observed and latent variables, whilst structural models describe the connection between latent variables.

In the current study, we used hierarchical SEM to examine the extent listeners’ physiology mediated the effect to which acoustic features underlying each musical expression shaped listeners’ pleasure based on their flattened twRDMs. On the first level, we first defined the latent variable ‘physiology’ to consist of HR, RVT, and pupil diameter as observed endogenous variables, as well as the latent variable ‘pleasure’ to consist of slider rating as the observed endogenous variable. Given that there were no significant differences between the lower noise-ceiling of listeners’ behavioural slider ratings and brightness and roll-off, we then defined the latent variable ‘acoustics’ comprising the two spectral features as observed exogenous variables. Next, we constructed three structural models: To test for direct effects, we set one model predicting pleasure from acoustics. To test for indirect effects, we defined one model to predict physiology from acoustics, and another to predict pleasure from physiology. The product of parameter estimates from the indirect effect models quantifies the mediation effect, and summing this product with the parameter estimate of the direct effect model gives us the total effect. Finally, to account for between-subject variability, the variance of RVT was modelled in the second-level. Though we considered other second-level covariance structures, this was the only structure that yielded a satisfactory model fit and was therefore selected.

The SEM model was estimated using maximum likelihood using lavaan 0.6-19 (107). Model fit was ensured by comparing our model versus a baseline model with no covariance between variables, which yielded a χ^2^-test p-value of 0.000, comparative fit index of 0.952, standardised root mean square residual of 0.03 (within-subject) and 0.00 (between-subject), and root mean square error of approximation (RMSEA) of 0.139, with p(H0: RMSEA>=0.08) = 1. We report p-values and bias-corrected confidence intervals of parameters estimated using bootstrapping with 5,000 samples with the manymome package (108).

### Testing for respiratory sinus arrhythmia

We used Granger causality to test for directed information transfer between HR and RVT timeseries in each trial for all subjects to assess the potential signs of respiratory sinus arrhythmia in our data (refer to Supplementary Figure 3). In brief, Granger causality tests whether the current value of a signal can be predicted by its own values earlier in time, as well as previous values of another signal. This was implemented using the VLTimeCausality package (109), which extends the Granger causality framework to allow for non-stationarity in the signal and relaxes the assumption of a fixed time-lag. Granger causality was assessed using an F-test with alpha = 0.05. Here, we say that there is directed information transfer from *X* to *Y* if *X* Granger-causes *Y* but not vice-versa, and we say that the test is inconclusive if the F-test is significant in none or both directions.

## Author contributions

Conceptualization: VKMC, SF, SS; Data curation: SS, TH, VKMC; Formal analysis: VKMC, TH; Funding acquisition: SF; Investigation: SS; Methodology: SS, SF, VKMC; Project administration: VKMC, SF; Resources: SF; Software: VKMC, TH; Supervision: SF, VKMC; Validation: VKMC; Visualization: VKMC, TH; Writing – original draft: VKMC, TH; Writing – review & editing: VKMC, TH, SS, SF

## Supplementary Information

**Supplementary Figure 1:**
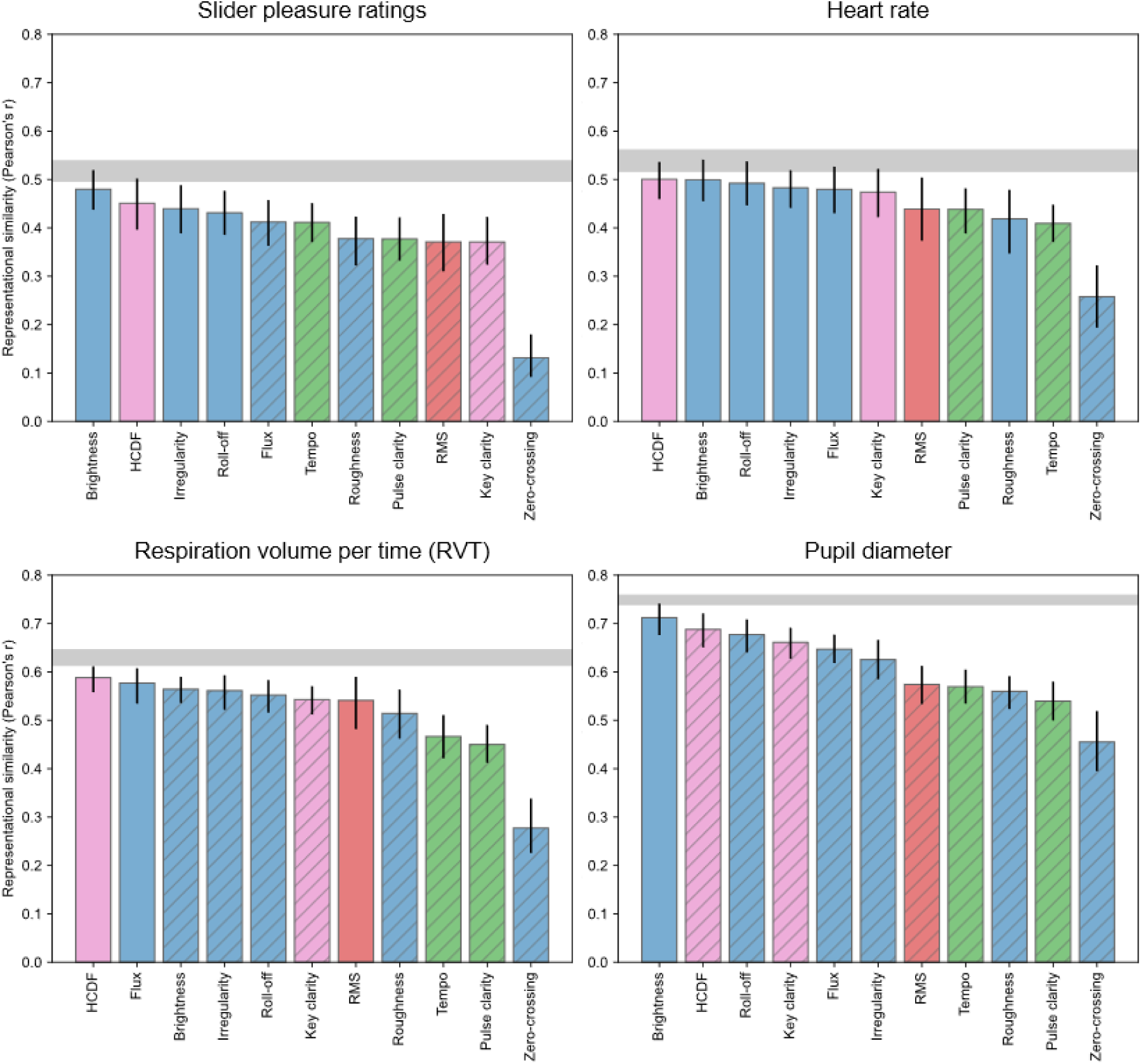
Representational similarity between acoustic features and behavioural or physiological responses with MIDI excluded. With MIDI excluded, HCDF and irregularity also did not differ significantly from the behavioural lower noise ceiling, suggesting additional candidate features that could capture changes in subjects’ behavioural responses. Likewise, brightness additionally did not differ significantly from cardiac and pupil noise ceilings. Refer to Figure 4 for legend.

**Supplementary Figure 2:**
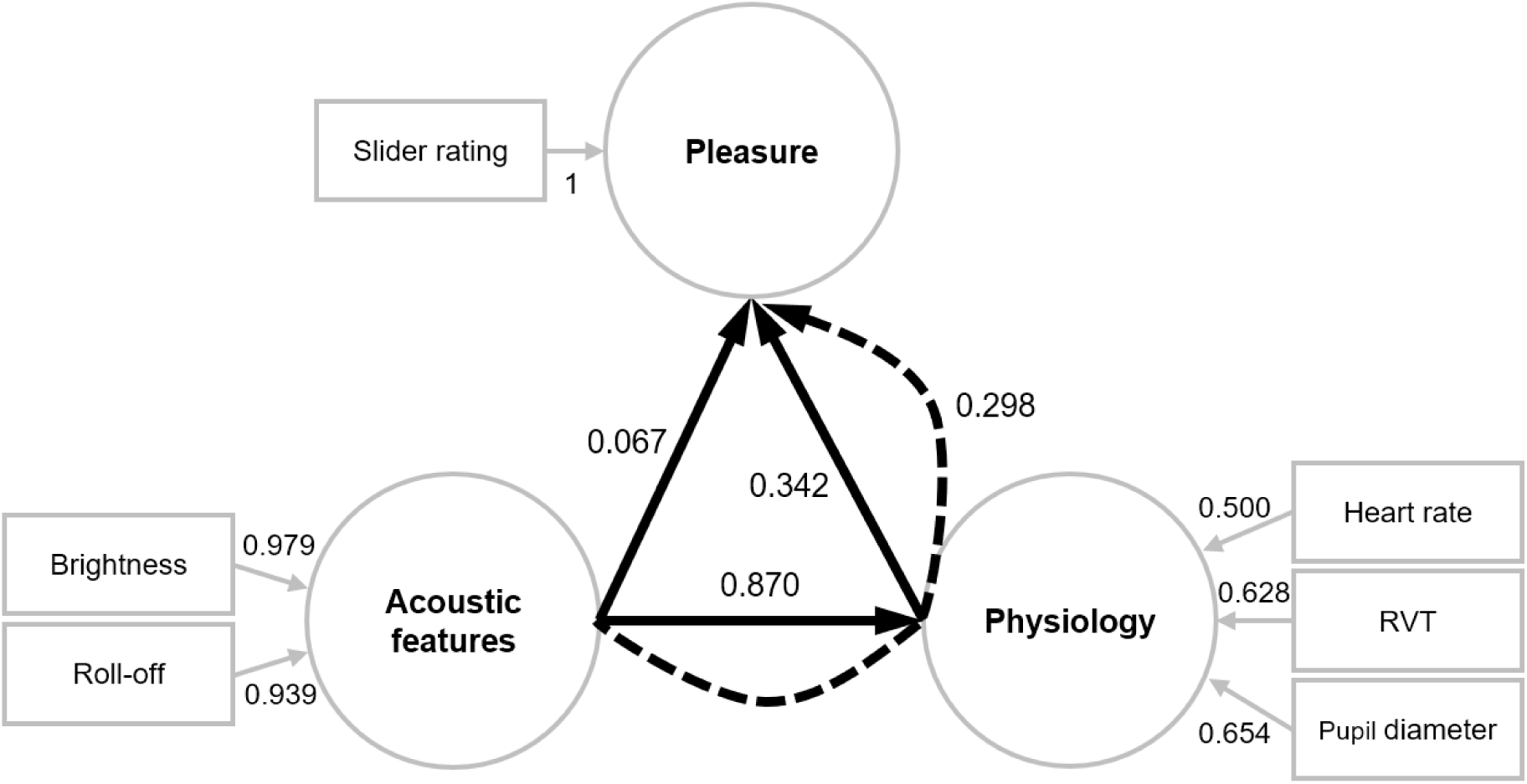
Structural equation modelling with MIDI excluded. Excluding MIDI from our structural equation model resulted in a larger direct effect of physiology on musical pleasure (0.342), as well as a larger indirect effect of acoustics on pleasure via physiology (0.298). These demonstrate the role of autonomic responses in shaping spectral acoustic-evoked changes in musical pleasure. Refer to Figure 5 for legend.

**Supplementary Figure 3:**
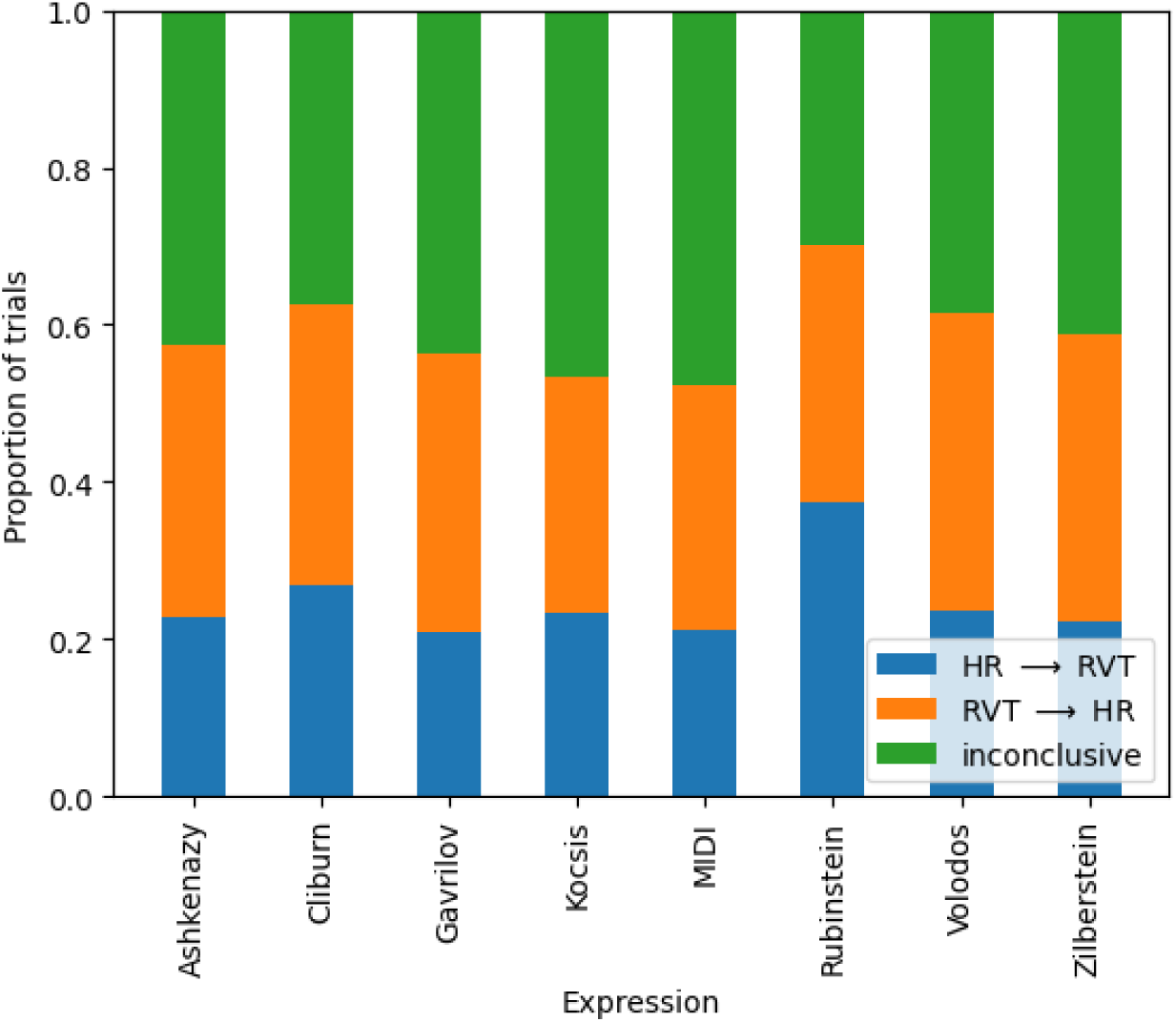
Minimal confound by respiratory sinus arrhythmia. Height of coloured bars indicate proportion of trials for each expression where earlier time lags in the heart rate (HR) timeseries could predict subsequent respiratory volume per time (RVT, blue), or vice versa (orange), or not (green). Only a small portion of trials (34.2%) indicated that earlier lags in RVT could predict heart rate, suggesting a minimal influence of respiration on cardiac activity in the current data.

**Supplementary Figure 4:**
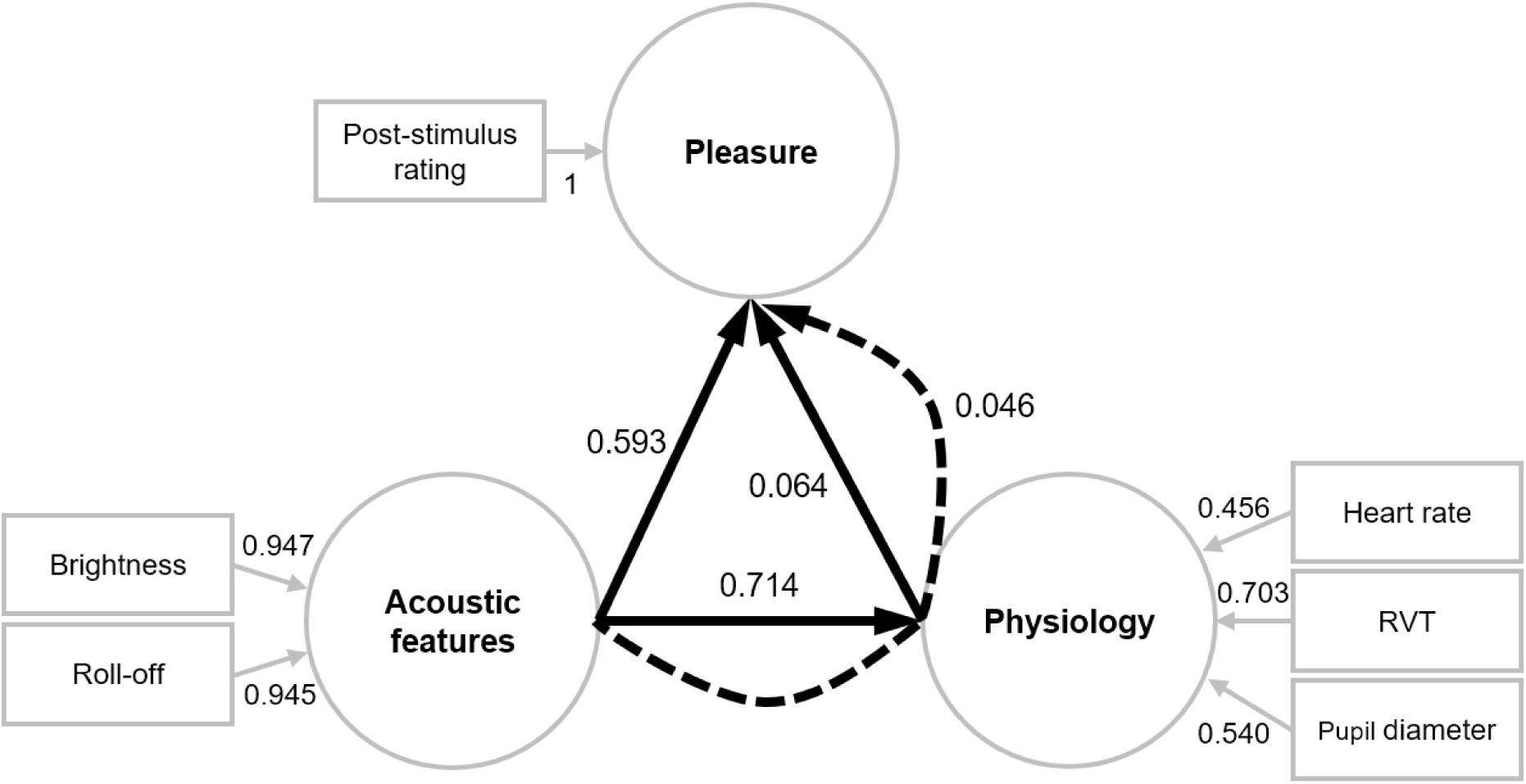
Structural equation modelling with post-stimulus rating. Although subjects’ post-stimulus pleasure ratings were highly correlated with the mean of their continuous slider pleasure ratings, we observed a far weaker direct effect between physiology and pleasure (0.064 vs 0.142), as well as the indirect effect of acoustics on pleasure via physiology (0.046 vs 0.101). Refer to Figure 5 for legend.

**Supplementary Figure 5:**
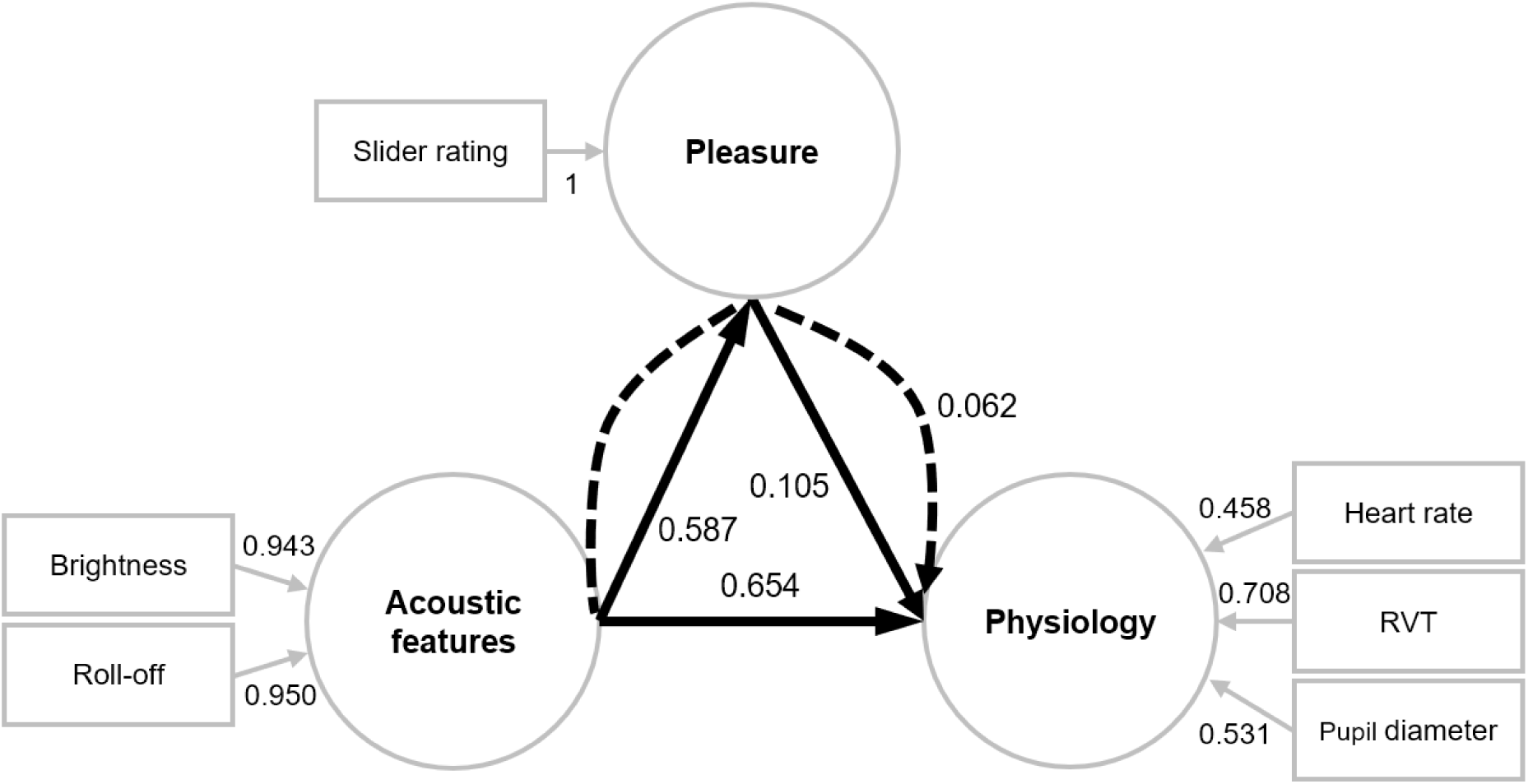
Reverse mediation testing. Interchanging pleasure and physiology as the mediation variable suggested a plausible mediation effect of pleasure on physiology, although this effect was 39% weaker than the mediation effect of physiology on pleasure (0.062 vs 0.101). Refer to Figure 5 for legend.

